# Impaired Hippocampal-cortical interactions during sleep and memory reactivation without consolidation in a mouse model of Alzheimer’s disease

**DOI:** 10.1101/828301

**Authors:** S. D. Benthem, I. Skelin, S. C. Moseley, J. R. Dixon, A. S. Melilli, L. Molina, B. L. McNaughton, A. A. Wilber

## Abstract

Spatial learning is impaired in preclinical Alzheimer’s disease (AD). We reported similar impairments in 3xTg-AD mice learning a *spatial reorientation task*. Memory reactivation during sleep is critical for learning related plasticity, and memory consolidation is correlated with hippocampal sharp wave ripple (SWR) density, cortical delta waves (DWs), and their temporal coupling - postulated as a physiological substrate of memory consolidation. Finally, hippocampal-cortical dyscoordination is prevalent in individuals with AD. Thus, we hypothesized impaired memory consolidation mechanisms in hippocampal-cortical networks could account for spatial memory deficits. We assessed sleep architecture, SWR/DW dynamics and memory reactivation in a mouse model of tauopathy and amyloidosis implanted with a recording array targeting isocortex and hippocampus. Mice underwent daily recording sessions of rest-task-rest while learning the *spatial reorientation task*. We assessed memory reactivation by matching activity patterns from the approach to the unmarked reward zone to patterns during slow wave sleep (SWS). AD mice had more SWS, but reduced SWR density. The increased SWS compensated for reduced SWR density so there was no reduction in SWR number. Conversely, DW density was not reduced so the number of DWs was increased. In control mice hippocampal SWR-cortical DW coupling was strengthened in post-task-sleep and was correlated with performance on the *spatial reorientation task* the following day. However, in AD mice SWR-DW coupling was reduced and not correlated with behavior, suggesting behavioral decoupling. Thus, reduced SWR-DW coupling may cause impaired learning in AD and may serve as a biomarker for early AD related changes.

**Significance Statement:** Understanding the relationship between network dynamics and cognition early in Alzheimer’s disease progression is critical for identifying therapeutic targets for earlier treatment. We assessed hippocampal-cortical interactions during sleep in AD mice as a potential cause of early spatial learning and memory deficits. We identified compensatory sleep changes in AD mice, that ameliorated some brain dysfunction. Despite the compensatory changes, impaired spatial navigation and impaired hippocampal–cortical (sharp wave ripple-delta wave) interactions were apparent in AD mice. In control but not AD mice hippocampal-cortical interactions were correlated with performance on the spatial task, the following day, suggesting a potential mechanism of impaired consolidation in AD mice. Thus, changes in hippocampal-cortical brain dynamics during sleep may underlie early memory deficits in AD.

## Introduction

Alzheimer’s disease (AD) is devastating for both individuals and society (McDade and Bateman, 2017). Individuals with AD have memory, cognitive and navigational impairments; in fact, getting lost and having impaired orientation in space is an early hallmark of AD (Allison et al., 2016; Henderson et al., 1989; Weintraub et al., 2012). Rodent models of AD are characterized by similar spatial navigation impairments (Attar et al., 2013; Liu et al., 2013; Marlatt et al., 2013). Spatial orientation and navigation involve hippocampal-parietal cortex (PC) interactions (Aguirre and D’Esposito, 1999; Byrne et al., 2007; Clark et al., 2018; Jarrard, 1993; Ji and Wilson, 2006; Kolb et al., 1994; Maingret et al., 2016; McNaughton et al., 1995; Morris et al., 1982; Nitz, 2012; Oess et al., 2017; Pai and Yang, 2013; Rogers and Kesner, 2006; Sherrill et al., 2013; Tu et al., 2017; Whitlock et al., 2012; Wilber et al., 2014; Wilber et al., 2017), and dysfunctional cortical-hippocampal interactions are a prominent feature in AD (Cacucci et al., 2008; Huijbers et al., 2014; Jacobs et al., 2012; Kunz et al., 2015; Morbelli et al., 2012; Mormino et al., 2012; Song et al., 2016; Wang et al., 2013), including abnormal communication between the PC and hippocampus (Jacobs et al., 2012; Kunz et al., 2015; Morbelli et al., 2012; Wang et al., 2013). These findings, make the hippocampal-PC network an ideal model for studying altered hippocampal-cortical interactions in AD (Cacucci et al., 2008; De Gennaro et al., 2017; Huijbers et al., 2014; Mormino et al., 2012; Song et al., 2016).

Memory consolidation involves cortico-hippocampal interactions, and changes in this network could be a contributing mechanism for AD-related memory impairments, particularly given evidence for altered cortical-hippocampal function in AD. The hippocampal formation is crucial to the storage of ‘episodic’ memories (memories for experiences that unfold in space and time), and for assisting the neocortex to extract generalized knowledge from these specific experiences (Eichenbaum, 2010). It has been suggested that the hippocampus generates a unique code reflecting the spatiotemporal context of experience that links together components of a given experience by producing interactions between modules throughout the neocortex, including the PC (Burke et al., 2005; Skelin et al., 2018). Hippocampus and PC exhibit coordinated replay during rest at both the single neuron (Qin et al., 1997; Wilber et al., 2017) and multi-neuronal (modular) level (Wilber et al., 2017). Memory reactivation has been proposed to be critical for neural changes underlying learning and memory (Ego-Stengel and Wilson, 2010; Girardeau et al., 2009; Jadhav et al., 2012; Staresina et al., 2013). In addition, mouse models of tauopathy and familial AD show changes in markers of memory replay. For example, the density and amplitude of hippocampal sharp wave ripples (SWRs) are reduced in these models (Ciupek et al., 2015; Gillespie et al., 2016; Witton et al., 2016). Additionally, later in AD progression, both animal models and humans show changes in slow-wave sleep (SWS) (De Gennaro et al., 2017; Mander et al., 2016; Van Erum et al., 2019). Finally, impaired navigation-related learning and memory in AD mice is largely a consequence of forgetting from one day to the next (Billings et al., 2005). Thus, memory consolidation, which involves cortico-hippocampal interactions is a potential mechanism for AD-related impairments. Specifically, pathology in the hippocampal-PC network may disrupt the physiological hippocampal-PC interactions that enable sleep-related memory replay and therefore disrupt the binding of aspects of a given experience (Braak and Braak, 1991; Khan et al., 2014; Mitchell et al., 2002). Here we examine hippocampal-PC brain dynamics during sleep following spatial learning behavioral sessions, to test the hypothesis that this system is disrupted in AD and may ultimately produce memory impairments in individuals with AD.

## Materials and Methods

Female 3xTg-AD (APP_Swe_, PS1_M146V_, and tau_P301L_) mice and age-matched NonTg mice (controls) from a similar background strain were group housed (2-4/cage) in 12:12 hour light-dark cycles until the experiment began. The triple transgenic mouse model, 3xTg-AD, expresses three major genes associated with familial AD, as well as the plaque and tangle pathology with a distribution pattern comparable to that observed in humans (Mesulam, 1999). Animals were originally obtained from Dr. Frank LaFerla (University of California Irvine) and bred in our vivarium. Mice were 6-7 months at beginning of experiment (n=5/genotype). All experimental procedures were carried out in accordance with the NIH Guide for the Care and Use of Laboratory Animals and approved by the University of California, Irvine and Florida State University Animal Care and Use Committees.

## METHODS DETAILS

### Pretraining

Mice were moved to single housing and water deprived to no less than 80% of starting weight. Then *alternation training* began where mice learned to shuttle back and forth along a linear track for a water reward that was delivered in the start-box only. At the end of the track was a black barrier in front (from the mouse’s perspective) of a black background. The starting position was moved to different locations, while the barrier remained fixed, allowing variation in track length. Starting positions were randomly selected, between 56-76cm from reward zone. All calculations were performed in pixels, and then converted to cm for visualization purposes. When mice met asymptote criteria (number of runs did not vary by more than ±6 on 3 of 4 days, or ran 50 or more times down the track in a single session), a date was scheduled for implantation of stimulating electrodes and recording array. Mice continued to run the task every other day leading up to implantation surgery, including the day before surgery.

### Surgical Procedure

*Two bipolar* stimulating electrodes were implanted unilaterally, targeting the left medial forebrain bundle (MFB; 1.9mm & 1.4mm posterior to bregma, ±0.8 mm lateral, 4.8 mm below dura). A 16-tetrode recording array (Chang et al., 2013) was implanted, targeting PC and hippocampus (2.2mm posterior to bregma, 2.0mm lateral). Animals recovered for one week, during which tetrodes targeting PC were moved down 31μm daily for the first three days, and then every other day, to prevent sticking. Tetrodes were not advanced beyond the lower border of PC, as determined based on inspection of LFP, and depth records. Tetrodes targeting hippocampus were turned down up to 124 microns daily for the first three days, and then 31 microns every other day, until tetrodes reached hippocampus, as indicated by depth records, as well as the characteristic hippocampal LFP.

### Stimulation parameters

After recovery, animals were placed in a custom 44 x 44 x 44cm black box with a nose poke port (Med Associates) in the left corner. Manual stimulation lasting 500ms was administered to shape the mice to use the nose poke port. Nose pokes were registered by a custom MATLAB program that automatically delivered stimulation. Once mice had been trained to use the nose poke port, stimulation parameters were varied (171-201Hz, 50-130μA current, electrode wire combination) to identify the settings that produced the highest response. No attempt was made to balance settings across genotype. However, response rate was compared across genotype to ensure that differences in reward strength were not likely to contribute to observed effects, and did not vary across genotype (t_(5)_=0.98, p=0.35). The same comparison was made for the two brain stimulation parameters that were adjusted, frequency and current. Frequency (t_(5)_=0.63, p=0.54) and current (t_(5)_=0.69, p=0.51) also did not vary across genotype.

### Spatial Reorientation Training

After optimal stimulation settings were identified, mice completed a spatial reorientation task previously described by (Stimmell et al., 2019) and similar to (Rosenzweig et al., 2003). Briefly, the *spatial orientation task* is identical to *alternation training*, except for the addition of an 8cm long goal zone in a fixed location in the room. For this task, if the mouse pauses in the real or virtual (for the virtual maze version described below) goal zone for a sufficient period, brain stimulation reward is delivered. The required duration of the pause in the goal zone gradually increases as the animal achieves asymptote criterion at each phase (0.5-2.5s in 0.5s increments). The real (or virtual) box and track are moved after each trial (sliding track or virtual teleportation), so the goal zone remains at a constant position within the real (or virtual) room; however, the goal zone can be at a variety of locations from the start box (40–110cm). The velocity profile during the approach to the reward zone (slowing) is used to assess performance (Rosenzweig et al., 2003; Stimmell et al., 2019). Asymptote criteria was ±15% correct trials on 3 out of 4 days. We found that the number of trials obtained on the real task were too low to reliably detect memory reactivation in NonTg mice therefore, all data reported here comes from the next (virtual) task for which mice run many more trials (378% increase).

### Virtual Maze

Upon completion of the spatial reorientation task, mice were trained on a virtual maze (VM; https://www.interphaser.com/, Molina et al., 2016) as described above. The only difference was the addition of an acclimation period for head fixation. Head fixation was performed manually for most mice (70%), except for a subset of mice that were able to tolerate head fixation utilizing a fixed head clamp. The tablet was coated with a thin layer of mineral oil and the mouse was allowed to run on a tablet, the mouse’s paws were sensed on the floor tablet similar to a finger swipe on a cell phone screen, which caused the virtual environment to move at the same rate as if the mouse was running across a real floor. Virtual movement was restricted to forward or backward only (the virtual environment could not turn). Three wall tablets (front and both sides) were synced to the floor tablet to display the view of the virtual room.

### Recording Procedures

An electrode interface board (EIB-72-QC-Small, EIB-36-16TT Neuralynx) was attached with a custom adapter to the recording array (Chang et al., 2013) with independently drivable tetrodes connected via a pair of unity-gain headstages (HS-36 Neuralynx) to the recording system (Digital Lynx SX Neuralynx). Tetrodes were referenced to a tetrode wire in the corpus callosum and advanced as needed, up to 62μm/day, while monitoring the audio and visual signal of the unit activity. Each daily recording session included a 50min sleep session, followed by 20min of task (real or virtual maze), followed by another 50min sleep session (Fig. 1A). Adjustments were made after a given day’s recording to allow stabilization overnight. Thresholded (adjusted prior to each session) spike waveforms were bandpass-filtered in 0.6 - 6kHz range and digitized at 32kHz. A continuous trace was simultaneously collected from one of the tetrode wires (bandpass-filtered in 0.1 - 1000Hz and digitized at 6400Hz) and referenced to an electrode in corpus callosum, was collected for processing as a local field potential (LFP). Mouse position was video-tracked using a colored dome of reflective tape for the real maze (or virtual position for the virtual maze), and on-line position information was used to trigger MFB stimulation rewards. Video-tracking or virtual position data was collected at 30Hz and co-registered with spikes, LFPs, and stimuli timestamps.

**Fig. 1.**
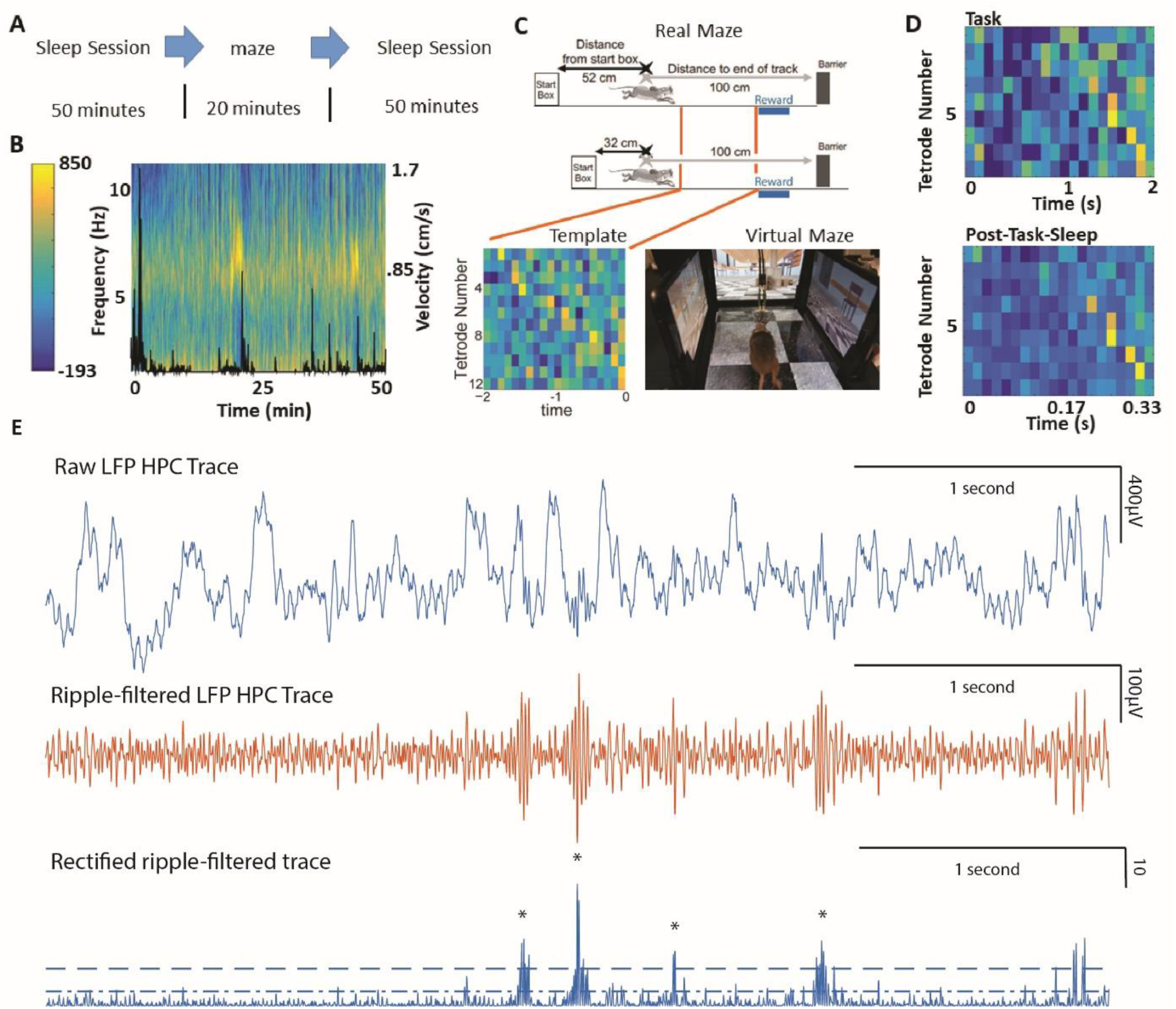
Daily recording sessions included sleep with state detection and a spatial reorientation task from which parietal cortex (PC) activity pattern templates were generated. **A.** All recording sessions started with a pre-task rest session, followed by the task (real or virtual maze), then a post-task rest session. Sleep sessions were 50min each and the spatial reorientation task or virtual reality task lasted 20 minutes for each daily recording session. **B**. Time frequency power spectrum for CA1 local field potential overlaid with movement velocity (black line) illustrates contrast between slow wave sleep (SWS; low velocity & low theta power), rapid eye movement sleep (REM; low velocity & high theta), and awake (high velocity & low theta). **C.** *Top.* The virtual maze task includes an unmarked rewarded location 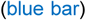 that is always fixed within the real or virtual room; however, the start box “moves” between trials. If only run in the long track configuration (*top*), the mouse could get position estimates from both self-motion (black) and real or virtual room cues 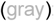; however, because there is a randomly selected set of track length configurations (e.g., *bottom*), room cues provide accurate position information, but self-motion information does not. Adapted from (Rosenzweig et al., 2003). *Bottom Left*. A template is constructed by averaging MUA across trials for each tetrode (rows) for the 2s period where the mouse approaches the reward zone on correct response trials. *Bottom Right.* We developed a mouse virtual maze (VM) system. Distal cues surround the virtual arena. There is a novel tablet-based control mechanism in which each mouse extremity is sensed just like a finger swipe on a phone. This results in many more trials (378% increase) on a 2D VM. **D.** A *task template* is constructed by averaging z-scored multi-unit activity (MUA) across trials for each tetrode (rows) for the last 2s window of the approach to the reward location (non-Tg mouse example; *top*). Next, the similarity is computed between this template and neural activity during a sliding window advanced one time bin at a time over SWS periods. Due to the compressed nature of memory reactivation, strong matches are only obtained when bin sizes are adjusted to apply a compression factor (e.g., for 6x *bottom* 0.33s sleep = 2s task). The top 100 matches were averaged to illustrate the strong correspondence between task template and activity patterns during post-task rest. Warm colors represent higher MUA Z-score for that time bin. **E**. SWRs were detected from the CA1 LFP (top) digitally filtered in the 75-300Hz (middle) and rectified (bottom). Events with peaks >6SD (dashed line) above the mean and duration 20-100ms were considered SWRs. The SWR duration included the contiguous periods surrounding the peak and exceeding 2SD (dotted line) above the mean.

Spike data were automatically overclustered using KlustaKwik and manually adjusted using a modified version of MClust (A.D. Redish). All spike waveforms with a shape, and across tetrode cluster signature suggesting that they were likely MUA and not noise, were manually selected and merged into a single MUA cluster. Thus, MUA clusters included both well-isolated single units and poorly isolated single units (http://klustakwik.sourceforge.net; Harris et al., 2000).

LFP analyses were performed using custom-written Matlab code (Mathworks, Nattick, MA) or Freely Moving Animal (FMA) Toolbox (http://fmatoolbox.sourceforge.net/). The LFP signal was collected at 6400Hz and subsequently resampled to 2000Hz for further analysis, using the Matlab *resample* function.

### Histology

At the conclusion of the experiment, marking lesions were made by applying a 5μA current for 10s between each tetrode wire and the tail. One week later mice were given an intraperitoneal injection of Euthasol and then transcardially perfused with 0.1*M* phosphate-buffered saline (PBS), followed by 4% paraformaldehyde (PFA) in 0.1*M* PBS. The whole head was post-fixed for 24h, to allow for easy identification of the tract representing location of tetrodes and MFB electrodes, and then the brain was removed and post-fixed for another 24h. Last, the brain was cryoprotected in 30% sucrose in 0.1*M* PBS. Frozen sections were cut coronally with a sliding microtome at a thickness of 40μm and split into 6 evenly spaced series.

#### 6E10

One series of sections was mounted on slides, incubated in 4% PFA for 4min, and then rinsed with Tris-buffered saline (TBS). Next, slides were soaked in 70% formic acid for 8 – 15min. After rinsing in TBS, slides were incubated in 0.1% Triton-X in TBS for 15min, followed by 0.1% Triton-X and 2% bovine serum albumin (BSA) in TBS for 30min. Sections were incubated with anti-β-amyloid 1-16 (mouse, clone 6E10, Biolegend) 1:1000 and anti-NeuN (polyclonal, rabbit, Milipore) in 0.1% Triton-X and 2% BSA in TBS for 2 days. After rinsing with TBS, slides were soaked in 0.1% Triton-X in TBS for 15min, followed again by 0.1% Triton-X and 2% BSA in TBS for 30min. Staining was visualized with anti-mouse-alexa-488 (1:1000) and anti-rabbit-alexa-594 (1:500) in 0.1% Triton-X and 2% BSA in TBS for 5-6h. Slides were coverslipped after being rinsed with TBS. Whole slides were imaged as described below, then the coverslip was removed and DAPI (0.01mg/ml) was added before the slides were re-coverslipped and reimaged. Except for β-Amyloid 1-16 as noted above, histology was performed on free-floating sections. Sections are permeabilized in 0.3% Triton-X and blocked in 3% Goat Serum in TBS, then incubated in primaries antibodies.

#### M78

Sections were rinsed twice in PBS, then blocked in PBS containing 0.3% Triton X and 3% goat serum. Sections were incubated for two days with 0.3μg/ml anti-MOC78 (monoclonal, rabbit, abcam 205341) in PBS containing 0.3% Triton-X and 0.02% sodium azide. Next, sections were rinsed with PBS and 0.3% Triton-X three times. Sections were then incubated in anti-rabbit-alexa-488 (goat, 1:500) in 0.3% Triton-X overnight. Sections were then rinsed with PBS and 0.3% Triton-X washes for 20min five times. Next, sections were incubated in anti-NeuN-Cy3 (polyclonal, rabbit, Milipore, ABN78C) 1:300 in PBS with 0.3% Triton-X and 0.02% sodium azide for 24h. Finally, sections were rinsed once with PBS containing 0.3% Triton-X, and twice with PBS. Sections were mounted with Vectashield containing DAPI and coverslipped and imaged.

#### M22

Sections were rinsed twice in PBS, then blocked in PBS containing 0.3% Triton X and 3% goat serum. Sections were incubated for two days with 0.5μg/ml anti-M22 (rabbit) in PBS containing 0.3% Triton-X and 0.02% sodium azide. Next, sections were rinsed three times in PBS with 0.3% Triton-X. Next, sections were incubated in anti-rabbit-alexa-488 (goat) 1:500 PBS with 0.3% Triton-X and 0.02% sodium azide overnight. Sections were then rinsed five times with PBS and 0.3% Triton-X. Sections were then incubated overnight in anti-NeuN-Cy3 (polyclonal, rabbit, Millipore ABN78C) 1:300 PBS with 0.3% Triton-X and 0.02% sodium azide overnight. Next, sections were rinsed once with PBS and 0.3% Triton-X, and then rinsed twice with PBS. Sections were mounted with Vectashield containing DAPI and coverslipped and imaged.

#### Thioflavin S

Anti-NeuN (1:1000), overnight, was followed by anti-rabbit-alexa-594 (1:500) for 5h. Sections were rinsed then immersed in a 1% Thioflavin S solution (Sigma) for 9min, rinsed in dH2O, destained in 70% Ethanol for 5min, rinsed in dH2O, and then transferred to TBS before mounting onto slides.

#### Phosphorylated tau

Incubation in anti-Phosphorylated tau (1:500, monoclonal, mouse, Thermo Scientific) with anti-NeuN (1:1000) overnight was followed by secondary antibodies, anti-mouse-alexa-488 (1:1000) and anti-rabbit-alexa-594 (1:500) respectively, for 6h. Sections were rinsed and mounted onto slides.

#### Parvalbumin

Sections were quench in 0.3% H_2_O_2_ in PBS for 25 minutes, then blocked in 5% goat serum in 0.5% Triton-X TBS for 90min. Primary antibody (mouse anti-parvalbumin; Sigma Aldrich) 1:2000 was added for 2 days, followed by a biotinylated goat anti-mouse antibody (Sigma Aldrich) 1:500 for 90min both in TBS with 0.5% Triton-X. Following this, A and B form the standard Vectastain ABC kit (Vector Laboratories) 1:500 in PBS was added for 1h. Staining was developed using a DAB (3,3′-Diaminobenzidine tetrahydrochloride hydrate; Sigma Aldrich) solution containing 0.05% DAB and 0.015% H_2_O_2_ in TBS. Sections were rinsed in PBS and mounted onto slides. After air drying, slides were dehydrated in increasing concentration of alcohol, cleared with Hemo-De and coverslip with Fisher Chemical Permount™ Mounting Medium.

### Image Acquisition

Whole slides were scanned using a scanning microscope at 40x magnification (NanoZoomer Digital Pathology RS Hamamatu) or 20x magnification (Zeiss Axioimager M2).

### Genotyping

We received homozygous 3xTg-AD mice from Dr. Frank LaFerla’s lab. We confirmed that all mice used in the experiment contained each transgene using conventional PCR. DNA was extracted from the tails of each mouse. Homozygosity was confirmed by cutting the PS1 PCR fragment with the *BstEII* restriction enzyme. Only the mutated human PS1 gene contains a *BstEII* cut site and will be cut. The absence of an uncut PCR product indicated that the mouse was indeed homozygous for the human PS1. The presence of overexpressed APP and Tau were also confirmed by PCR. The previously published primers were used for amplifying the PS1 transgene, APP and Tau (Stimmell et al., 2019).

## QUANTIFICATION AND STATISTICAL ANALYSIS

### Sleep and LFP Analyses

First, still periods were extracted from the rest sessions as described previously (Euston et al., 2007; Wilber et al., 2017). The raw position data from each video frame was smoothed by convolution of both x and y position data with a normalized Gaussian function (standard deviation of 120 video frames). After smoothing, the instantaneous velocity was found by taking the difference in position between successive video frames. An epoch during which the velocity dropped below 0.78pixels/s (~0.19cm/s) for more than 2min was considered a motionlessness period. All analyses of rest sessions were limited to these motionless periods. Slow wave sleep (SWS) and rapid eye movement (REM) sleep were distinguished using automatic K-means clustering of the theta/delta power ratio extracted from the CA1 pyramidal layer LFP recorded during the ‘stillness’ periods, as is commonly done in a variety of major memory reactivation labs (Fig. 1B; Buzsáki et al., 2003; Girardeau et al., 2009; Grosmark et al., 2012; Lansink et al., 2008; Mizuseki et al., 2011; Mizuseki et al., 2009; Sirota et al., 2008; Stan Leung, 1998). Only slow wave sleep periods were included in the analysis. Delta wave troughs (DWT), which correspond to cortical down states (Battaglia et al., 2004), were detected by digitally filtering the LFP trace from a representative cortical electrode in the 0.5-4Hz range and detecting the peaks in the inverted signal that exceeded a mean+1.5SD threshold. SWRs were detected from the CA1 LFP digitally filtered in the 75-300Hz. Events with peaks >6SD above the mean and duration 20-100ms were considered SWRs. The SWR duration included the contiguous periods surrounding the peak and exceeding 2SD above the mean. The SWR detection accuracy was visually validated on the subset of each analyzed dataset. This visual validation process caused us to adjust the SWR detection criteria for two (one NonTg, one 3xTg-AD) mice. For these mice events with peaks >7SD were used instead.

### Template Matching

We performed *template matching* analysis as was done previously to show the simultaneous reactivation of isolated single cell ensembles in the medial prefrontal cortex (Euston et al., 2007; Louie and Wilson, 2001) and that we applied more recently to MUA in PC (Wilber et al., 2017). The criteria for the inclusion of a dataset in the template matching analysis was at least 20min of stillness and 600DWTs during both pre- and post-task-sleep, and at least 10 complete trials during the task period. Template matrices (number of tetrodes x number of time bins; Fig. 1C) were generated from the trial-averaged multiunit activity (MUA template) extracted from a 2s/trial windows that preceded the arrival at the reward site and binned to 100ms bins. The time window was chosen based on evidence that reactivation of the hippocampal activity patterns is more prominent for the task phase immediately preceding the reward (Diba and Buzsaki, 2007; Foster and Wilson, 2006; McNamara et al., 2014; Singer and Frank, 2009; Wilber et al., 2017). The time bin choice was based on previous reports of template matching using isolated single unit activity (Euston et al., 2007; Johnson et al., 2010), where a 100ms bin was deemed optimal for capturing task-related neuronal dynamics and identical to what we used previously (Wilber et al., 2017). In order to eliminate tetrodes with sparse and/or poorly approach-modulated activity, only the tetrodes with average reward approach period MUA >1Hz and reward approach period binned spike train coefficient of variation >0.25 were included in the template. Coefficient of variation was calculated over twenty 100ms time bins. Only the templates containing ≥6 tetrodes or single cells in PC were retained. To eliminate the influence of signal amplitude variability between tetrodes, the binned signal was z-scored for each tetrode separately.

Due to the time-compressed nature of re-activation reported previously for single-cell (Euston et al., 2007; Peyrache et al., 2009; Wilson and McNaughton, 1994) and MUA (Wilber et al., 2017) memory reactivation studies, we performed template matching for several evenly spaced compression factors: ‘no-compression’, 4x, 6x, 8x, and 10x. All of the template matching analyses were limited to SWS periods. Signal from the sleep periods was processed in the same way as for behavioral templates, except that the bin size was adjusted according to the compression factor (bin size=100ms/compression factor; e.g. for the 4x compression, the sleep bin size=25ms), to capture the compressed nature of neural reactivation during sleep.

To test the matching of a given template and the pattern of activity during sleep *(matching significance)*, we employed the approach identical to what we used previously in rats (Wilber et al., 2017). First, each template was shuffled repeatedly to generate 100 *shuffled templates*. The shuffling procedure consisted of randomly permuting the position of each column in the template (*population vector*), preserving the overall activity levels and instantaneous correlations between the signals on different tetrodes, but scrambling the sequential patterns. A Pearson correlation coefficient was calculated between each template and the series of *candidate matches*, generated by sliding the template-size window over the sleep epoch. This resulted in a matrix of Pearson correlation coefficients r, where the element r^i,j^ corresponded to the correlation coefficient between the i-th template and j-th candidate match. The correlation matrices were z-scored across individual time bins (columns), and the resulting z-score values reflected the template similarity to the corresponding sleep segment at given time step, relative to the distribution that included the original and 100 shuffled templates. Z-score values above 3 were considered *matches*.

For comparison of template matching between the pre- and post-task-sleep, *match percentage* was obtained by dividing the number of matches by the number of SWS time bins for each sleep epoch. For the comparisons between the original and shuffled templates, means and distributions of z-score values obtained from the original and shuffled templates were compared within epoch. For the DWT-triggered, the original template z-score traces +/-1s around each DWT most-negative point time were averaged, obtaining the event-triggered averaged z-score for a given sleep epoch. The peri-event reactivation strength was defined as the maximum averaged event-triggered z-score value within +/− 0.7s window surrounding detected event. To assess hippocampal-PC interactions, cross-correlations were performed between DWTs in PC, and hippocampal SWRs, both for *post-task-* and *pre-task-* sleep. SWR counts with bin size 16.67ms were centered on the DWT and then z-scored, and z-scores were compared between *pre-task-* and *post-task-* sleep.

## Results

### Memory Reactivation Characteristics in NonTg Mice

First, we tested the degree of temporal synchrony between the hippocampal and PC memory replay markers (SWRs and DWTs) in NonTg mice (Fig. 2A). Cortical replay tends to occur during the transitions between down- and up-states, and short lasting down states correspond to the DWT (Peyrache et al., 2009). Further, template matching in PC is stronger during memory replay events in the hippocampus (SWRs), even after accounting for the influence of simultaneous population memory replay events in PC (i.e., delta-waves; Wilber et al., 2017). Therefore, we assessed memory replay interactions across regions by cross-correlating a replay marker in PC, the DWTs, with replay marker from hippocampus, SWRs. We found that during SWS periods, NonTg mice had increased occurrence of SWRs just before DWTs (−117 ms on Fig. 2B) in *post-task-*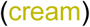 versus *pre-task-* (black) sleep. Next, we captured this effect by measuring the z-scored peak in the cross-correlation for *pre-task-* compared to *post-task-* sleep in NonTg data sets that met inclusion criteria (≥10 trials on VM, ≥20 minutes of still per sleep session, & ≥600 DWT per sleep session) were divided into equally sized blocks of 3 days (n=24 data sets). We found that during days 1-3, SWR by DWT interactions were strengthened during *post-task-* sleep, but this increase was not statistically significant (Fig. 2B; paired t-test for peak in *post-task-* versus *pre-task-* sleep: t_(8)_=-1.31, p=0.22). Similarly, during days 4-6, SWR by DWT interactions were significantly strengthened during *post-task-* sleep (t_(6)_=-3.80, p<0.05). However, during days 7-9, SWR by DWT interactions no longer robustly increased in post-task sleep and instead there was now a stronger negative than positive relationship between DWTs and SWRs (Fig. 2B).

**Fig. 2.**
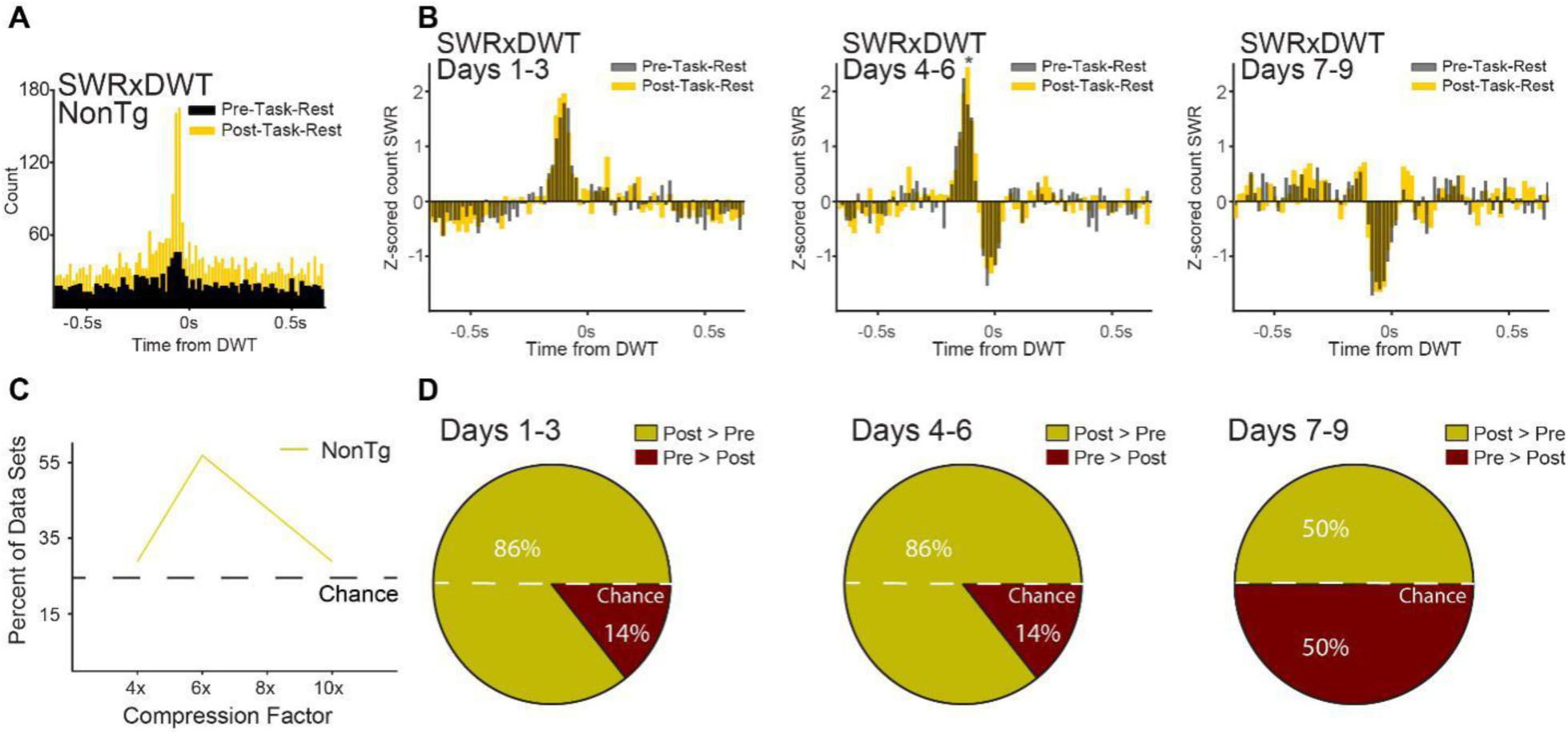
Memory replay measures suggest a decreased reactivation strength after 6 days of virtual maze (VM) training. **A.** Example cross-correlation between hippocampal (HPC) sharp-wave ripples (SWR) and parietal cortex (PC) delta wave troughs (DWTs) shows that SWRs tend to precede DWTs and this relationship is strengthened in *post-task-rest* 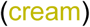. **B.** Average z-scored cross-correlation between hippocampal SWRs and PC DWTs for all data sets in NonTg animals that met the inclusion criteria in days 1-3, 4-6, and 7-9 of the VM task. The temporal correlation between HPC SWR and DWT decreases sharply after the first 6 days of VM training. The peak of the cross-correlation exceeded 1SD in the positive direction for days 1-3 & 4-6, but not 7-9 and was significantly greater in post-task sleep than pre-task sleep for days 4-6 (*p<0.05). **C.** The percent of data sets for which template matches are greater in *post-task-sleep* and also greater than non-compressed data for NonTg 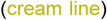 mice indicates a peak in template matches for the 6x compression factor**. D.** The proportion of data sets in which the proportion of template matches was larger in *post-task-sleep* for all NonTg mouse data sets that met inclusion criteria for days 1-3, 4-6, and 7-9 of VM training, Template matching in parietal cortex (PC) parallel the SWR-DWT coupling strength time course and also falls off sharply in days 7-9.

Next, we looked for evidence that NonTg mice have memory replay in the PC that is detectable using template matching, as we found in rats (Wilber et al., 2017). Note, we operationally define memory reactivation or replay as matching activity patterns from a *task* to *post-task-sleep* and we are not directly measuring ‘memory’ per se. Cortical memory replay is temporally compressed (Fig 2; Euston et al., 2007; Wilber et al., 2017). Thus, a control is to compare template matching for temporally compressed versus non-compressed templates. Therefore, we assessed template matching for a range of evenly spaced compression factors (‘non-compressed’, 4x, 6x, 8x, and 10x; as in our previous paper Wilber et al., 2017). We found template matching was strongest when a compression factor of 6x was applied (i.e., sequences of activity are reactivated 6x faster than during behavior; Fig. 2C), consistent with previous findings of temporally compressed memory replay for both single neuron and multi-neuronal activity patterns (Battaglia et al., 2004; Diba and Buzsaki, 2007; Euston et al., 2007; Peyrache et al., 2009; Wilber et al., 2017).

Finally, we assessed potential influences of time or training phase on memory reactivation strength. To do this we focused on the first 9 days of the VM task, the same time course as for the HPC-SWR cross correlation analyses for data sets that met the inclusion criteria (n=20 data sets; same as for the SWR-DWT cross-correlation analyses but with one additional criteria of ≥6 tetrodes in PC). During days 1-3 and 4-6 of the VM task, identical to the observation for DW-SWR cross-correlations, for a majority of the data sets (86%) there was an increase in proportion of strong template matches in post-task sleep during SWS (Fig. 2D). However, during Days 7-9, the proportion of data sets showing stronger template matching in post-task sleep falls to chance (50%). Thus, for the VM task in mice, PC memory replay is strong during the first 6 days of training, then falls off sharply.

### Impaired Spatial Learning and Memory

Next, we looked for impaired virtual spatial re-orientation performance to see if similar deficits occurred on the VM to our published data from the real maze (n=5 mice/genotype; Stimmell et al., 2019). As in our previous paper, 6-month 3xTg-AD mice have low levels of pathology with only intracellular staining for pTau, 6e10, M22, and M78 and no extracellular plaques or tangles, e.g., ThioS (Stimmell et al., 2019).Because mice are teleported to a random start location and the goal zone is unmarked, mice must use distant maze cues to get oriented in the maze space in order to locate the hidden reward zone and receive a reward (Fig. 1C). First, we assessed NonTg performance on the VM. Velocity for each day of the task was z-scored, to control for variations in running speed across animals. In order to quantify this apparent slowing in the reward zone in NonTg animals, we compared velocity in the reward zone with velocity immediately prior to the reward zone. NonTg mice slowed in the reward zone in days 1-3, as well as days 4-6 of the VM task (Fig. 3 *Top*; Fs_(1)_≥8.33, ps≤0.001). This slowing indicates the ability to use distal cues from the virtual room around the maze to locate the reward zone. When we performed the same analysis for days 7-9, NonTg mice continued to slow in the reward zone despite the increasing task difficulty as reward delays ranged from 1s to 1.5s by this point in training and testing (F_(1)_=28.71, p<0.0001). Next, we compared NonTg and 3xTg-AD mice and found that NonTg mice slowed more in the hidden virtual goal zone than 3xTg-AD mice (Fig. 3 *Bottom*; F_(1, 6)_=7.69, *p<*0.05). There was no effect of day or interaction between day and genotype (Fs_(5, 30)_<1.75, p>0.15). Thus, NonTg mice were better at learning to use distant maze cues to locate the virtual goal zone than 3xTg-AD mice. We performed the same assessment on days 7-9 of the VM task, comparing NonTg and 3xTg-AD mice and found that 3xTg-AD mice no longer performed worse than nonTg mice (F_1,6)_=3.30, p=0.12). Therefore, given that both template matching in PC and also interactions with hippocampus dropped dramatically after the first 6 days of VM training, and that impairments in 3xTg-AD mice were only present for the first 6 days, we limited our assessment of memory reactivation to the first 6 days of VM training and testing.

**Fig. 3.**
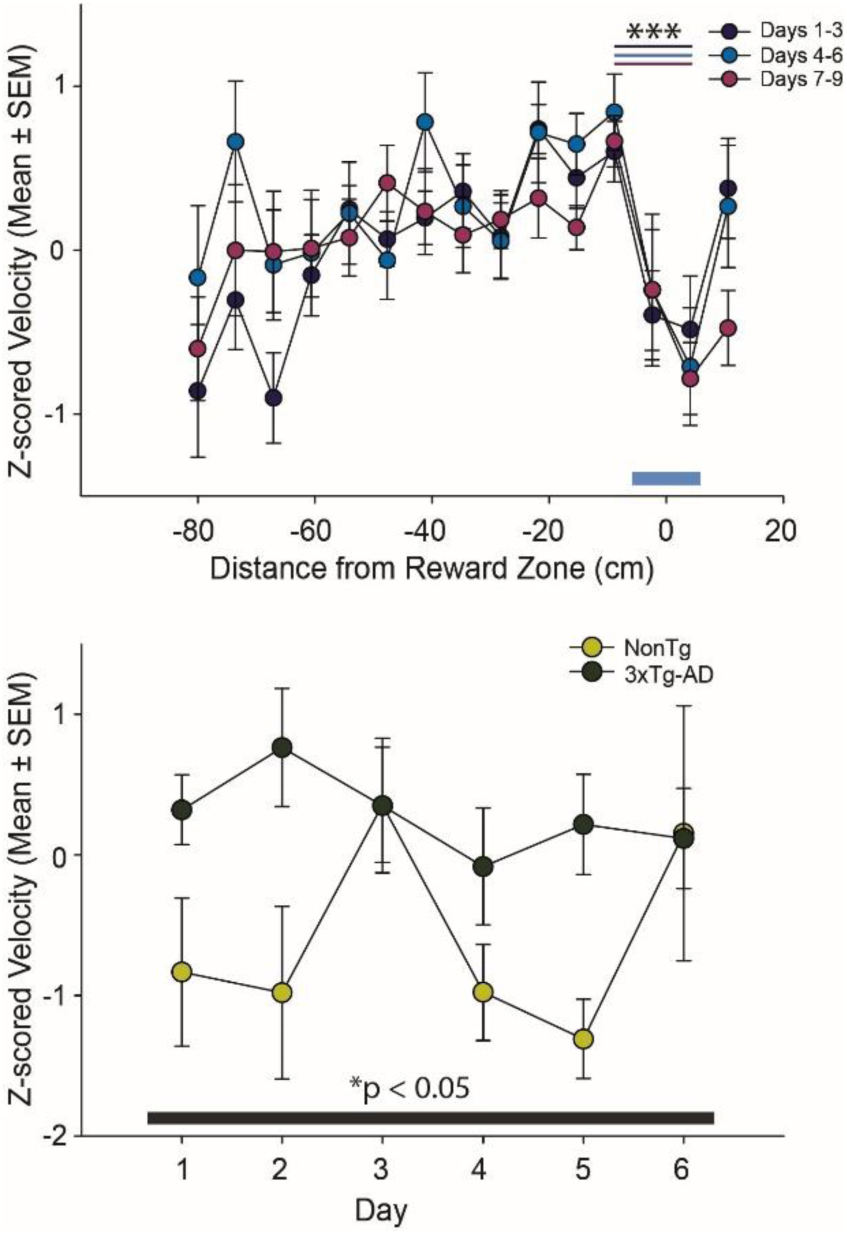
Impaired Spatial Learning and Memory on the Virtual Spatial Reorientation Maze. *Top*. Average Z-scored velocity Mean±SEM as a function of distance from reward zone 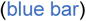 for NonTg mice for days 1-3 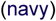, days 4-6 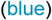 and days 7-9 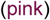 reward delays. NonTg slowed in reward zone, indicating knowledge of reward zone location based on maze cues in the virtual room (F_(1, 22)_≥8.32, p≤0.001). *Bottom*. Velocity in reward zone for NonTg 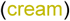 and 3xTg-AD 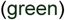 mice during first 6 days of behavior. 3xTg-AD mice slowed significantly less in reward zone regardless of day (F_(1, 6)_=7.69, p≤0.05).

### Compensatory Sleep Changes

Effects on sleep quality have been reported in AD (Roh et al., 2012a); however, those reports focused on later stages of disease progression (e.g., post-plaque formation). The 6-month 3xTg-AD female mice examined here have intracellular phosphorylated Tau and Aβ (Stimmell et al., 2019) but no extracellular tangles or plaques, so sleep quality in 3xTg-AD mice at this stage of disease progression isunknown (e.g., time still, proportion still time in SWS, proportion still in rapid eye movement sleep). Exclusion criteria were the same as for the cross-correlation analysis previously described, but based on our timecourse analysis we limited all further analyses to the first 6 days of behavior, resulting in 16 NonTg data sets and 11 3xTg-AD data sets. Surprisingly, we found that NonTg mice were motionless less than 3xTg-AD mice suggesting that 3xTg-AD mice actually had higher sleep efficiency than NonTg controls (Fig. 4 *Top Left*; effect of genotype: F_(1, 25)_=12.83, *p*<0.01 but not pre-versus post-sleep phase or interaction between genotype and sleep phase: Fs_(1, 25)_<0.87, *ps>*0.36). NonTg mice also had shorter average uninterrupted duration for each SWS episode (significant effect of genotype: F_(1, 25)_=14.29, p<0.01, but no sleep phase or interaction: Fs_(1, 25)_<1.12, ps>0.30). Interruptions to SWS bouts could be caused by either entering REM sleep or waking, so we next assessed REM sleep. We found that REM sleep proportion decreased during *post-task-sleep* (F_(1, 25)_=4.67, p<0.05) regardless of genotype (no effect of genotype or genotype by sleep phase interaction Fs_(1, 25)_<2.50, ps>0.14). Similarly, the number of REM bouts decreased in *post-task-sleep* (**Supplementary Figure 1**; F_(1,25)_=4.52, p<.05) regardless of genotype (no main effect of genotype or genotype by session interaction; Fs_(1, 25)_<0.32, ps>0.57). This decrease in REM sleep corresponded to an increase in SWS from pre-task-to post-task-sleep (F_(1, 25)_=6.33, p<0.05) regardless of genotype (no main effect of genotype or genotype by session interaction; Fs_(1, 25)_<2.69, ps>0.11). Thus, SWS increased after the task regardless of genotype and NonTg mice slept less than AD mice.

**Fig. 4.**
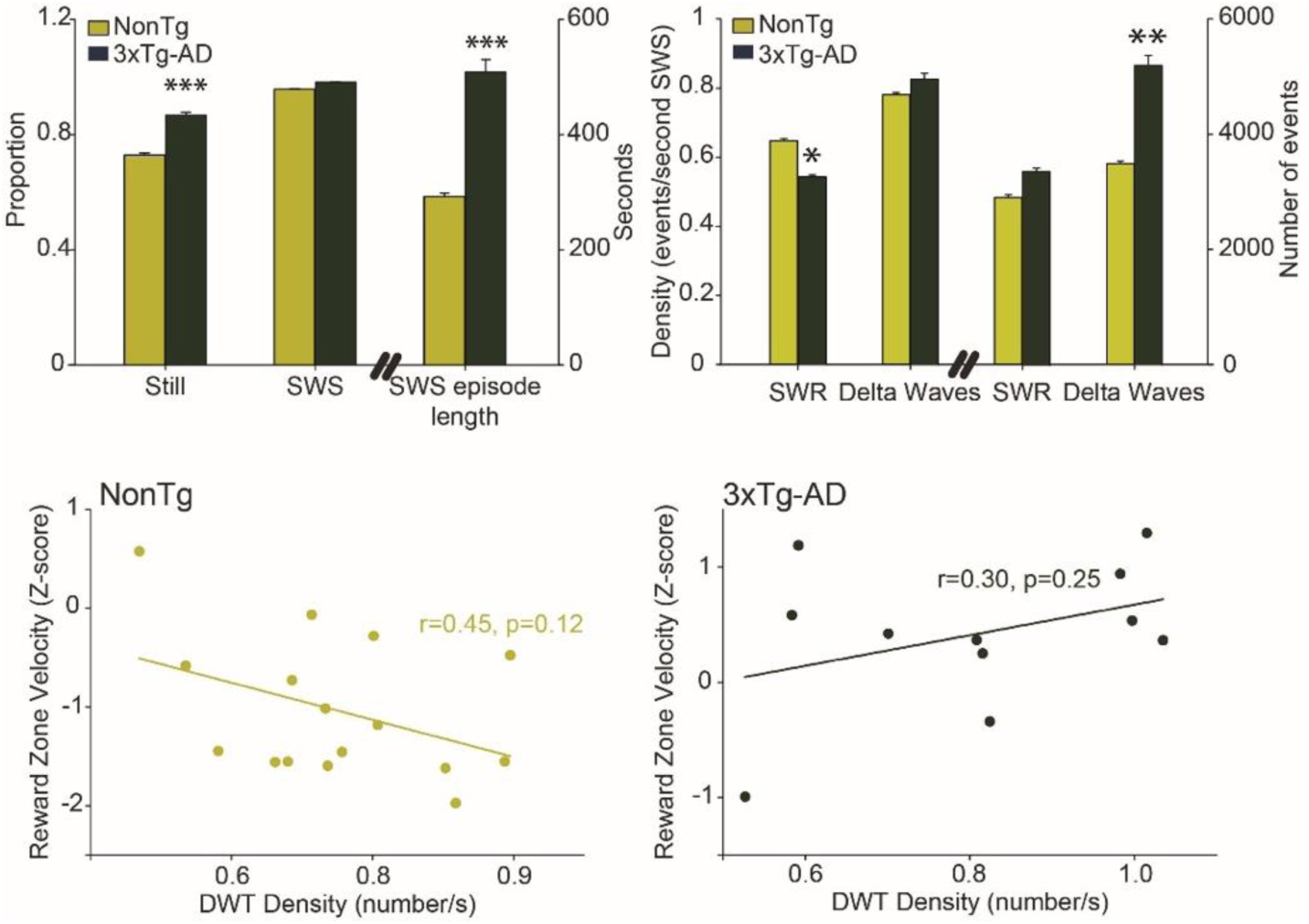
Sleep changes in AD mice compensate for a decreased sharp wave ripple (SWR) density. *Top Left*. Mean (±SEM) proportion of stillness and slow wave sleep (SWS), and SWS bout length for NonTg 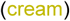 and 3xTg-AD mice (green). Transgenic mice spent a greater proportion of the sleep session still and had increased average length of SWS bouts. *Top Right*. Memory reactivation markers were examined: parietal cortex (PC) delta waves and hippocampal sharp wave ripples (SWRs). Density (*left*) and total number (*right*) of SWRs and DWTs from stillness time periods. Transgenic mice had decreased SWR density and increased DWT number. Repeated measures ANOVAs did not show significant effects of sleep phase (pre-task and post-task sleep) or interaction with genotype, so pre- and post-task-sleep were combined here for illustrative simplicity. *Bottom.* Higher DWT rate was not a significant predictor of behavioral performance (z-scored velocity in the reward zone) the following day for NonTg (*Left*) and 3xTg-AD mice (*Right*); however, this relationship was negative for NonTg mice (higher DWT was related to better performance) but was positive for 3xTg-AD mice (higher DWT density was not related to better performance). * p ≤ 0.05 ** p≤0.01. *** p ≤ 0.001.

Next, we individually assessed markers of memory reactivation, DWT in PC and SWRs in hippocampus (Fig. 4 *Top Right*). Since 3xTg-AD were still for a higher proportion of the sleep sessions and had increased SWS episodes compared to NonTg mice, we assessed both the density and the total number of these events. The density of SWR was decreased in 3xTg-AD mice, relative to NonTgs (F_(1, 25)_=5.54, p<0.05, no group or session interaction; Fs_(1, 25)_<1.83, ps>0.18). However, the total number of events was not different between groups, with no main effect of group or interactions (Fs_(1, 25)_<2.72, ps>0.11). The density of DWT was equivalent between NonTg and 3xTg-AD (F_(1, 25)_=0.23, p=0.63, no group or session interactions; Fs_(1, 25)_<0.89, ps>0.35), so the total number of DWT was increased in 3xTg-AD mice (F_(1, 25)_=9.83, p<0.01, no group or session interaction; Fs_(1, 25)_<1.5, p>0.22).

Next, we assessed the relationship between the density and number of SWRs and DWTs during *post-task-sleep* and behavioral performance the following day. Total number and density of SWRs was not a significant predictor of performance on the VM the following day for either NonTg or 3xTg-AD mice (rs<0.40, ps>0.22). Total number of DWTs also did not predict performance on the subsequent day for either group (rs<0.17, ps>0.14). Similarly, DWT density was not a significant predictor of performance on the VM for NonTg mice the following day (Fig. 4, *Bottom left;* r=-0.41, p=0.12). When we performed the same comparison for 3xTg-AD mice, DWT density again did not significantly predict performance on the VM the subsequent day but the direction of relationship between higher DWT density and slowing in the reward zone was reversed (i.e., slope of the regression line was positive; Fig. 4, *Bottom right;* r=-0.38, p=0.25). Thus, post-training sleep DWT and SWR density *alone* were not significant predictors of performance on the VM the following day.

### Impaired Hippocampal-PC Interactions

Next, we assessed HPC-PC interactions to see if 3xTg-AD mice had changes in network dynamics that could reflect miscoordination of memory replay (n=11 3xTg-AD data sets met the inclusion criteria described above for nonTg mice). We assessed hippocampal-PC coupling by cross-correlating a replay marker in PC, the DWT, with one from hippocampus, SWRs, as described above for nonTg mice. First, we assessed the proportion of data sets with a larger *post-task* verses *pre-task--sleep* peak in the SWR by DWT correlation. In NonTg mice, a significantly larger proportion (Fig. 5 *Top Left*) of data sets (75%) showed an increase in the SWR by DWT correlation, compared to 3xTg-AD mice (36%; Fig. 5 *Top Right*, χ^2^_(1)_ =4.03, p≤0.05). This suggests that coordination of memory replay events was impaired in AD mice. Next, we assessed the relative timing of these two events. We found that for SWS periods, NonTg mice had increased occurrence of SWRs just before DWTs (−117ms, peak left of lag zero on Fig. 5 *Middle Left*) in *post-task-*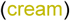 versus *pre-task-* (black) sleep (t_(15)_=2.92, p<0.05). However, when we performed the same assessment on 3xTg-AD mice, we found that while SWRs still tended to precede DWTs, this relationship was not strengthened during post task sleep (Fig. 5 *Middle Right*, t_(10)_=0.50, p=0.63). In fact, the mean cross-correlation peak from *pre-task*-sleep was taller than the *post-task*-sleep peak. This also suggests that cross-region coordination is impaired in the 3xTg-AD mice.

**Fig. 5.**
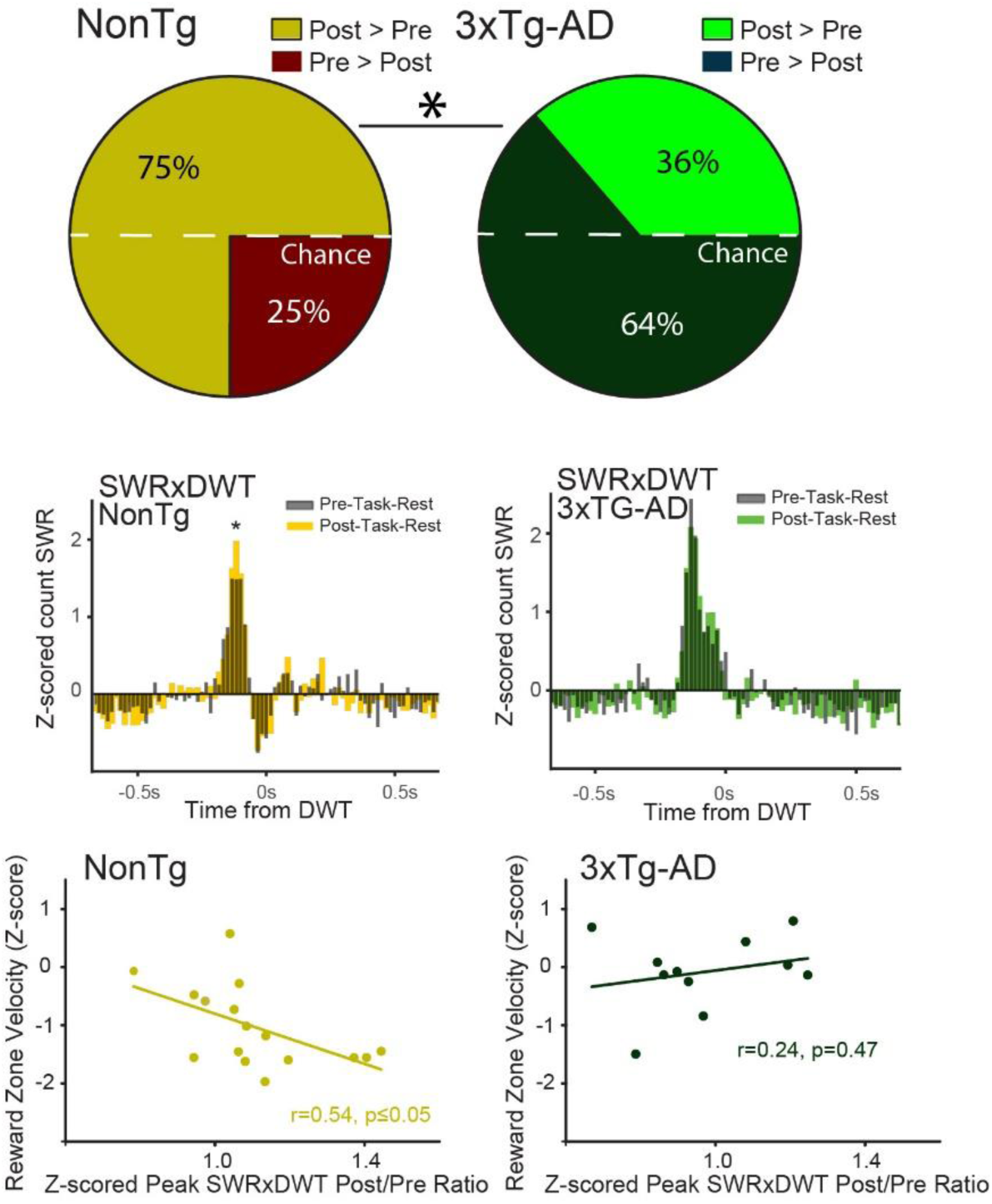
Parietal-hippocampal network coordination is impaired in 3xTg-AD mice and decoupled from behavior. *Top.* The proportion of data sets in which the cross-correlation peak was larger in *post-task-sleep* for all NonTg mouse data sets 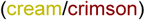 and all 3xTg-AD mouse data sets that met inclusion criteria (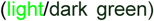/dark green) shows that HPC-PC reactivation event coupling is reduced in 3xTg-AD mice for *post-task* relative to *pre-task sleep. Middle Left.* Mean Z-scored cross-correlation between HPC SWR and PC DWT for NonTg data sets. SWRs tend to precede DWT by about 117ms, and that this relationship is strengthened in *post-task* 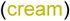 versus *pre-task-sleep* (black; *p<0.05). *Middle Right.* In 3xTg-AD mice, with SWR preceding DWT by an average by a similar lag as NonTg mice, about 133ms; however, the correlation between SWRs and DWTs is not strengthened in *post-task* (green) versus *pre-task* (grey)-sleep. *Bottom Left.* In NonTg mice strengthening of DWT-SWR coupling (Z-scored peak SWRxDWT post/pre ratio) is associated with better behavioral performance the following day (slowing behavior in reward zone; r=0.54, p<0.05). *Bottom Right.* In 3xTg-AD mice, strengthening of DWT-SWR coupling is not predictive of better behavioral performance the following day, and even shows a slightly positive relationship.

Finally, we assessed the relationship between SWR by DWT correlations and performance on the VM. We calculated z-scored peak ratio by dividing the z-scored peak from *post-task-sleep* by the z-scored peak from *pre-task-sleep* and compared this value to performance on the maze the subsequent day. In NonTg-AD mice, this ratio predicted performance on the VM the next day (Fig. 5 *Bottom Left*, r=0.54, p<0.05). However, 3xTg-AD mice did not have a significant relationship between this ratio and subsequent performance (Fig. 5 *Bottom Right*, r=0.24, p=0.47). Interestingly, as with DWT correlations, the direction of the relationship (i.e., slope of the regression line) was reversed in 3xTg-AD mice. As an additional control, we assessed the relationship between this ratio and performance on the task the same day. For both groups of mice, there was not a significant relationship between performance on the maze just prior to the sleep session that was assessed for SWRs by DWT correlations (rs<0.47, ps>0.14). This suggests that SWR by DWT correlations may represent plasticity that produces better performance the following day and that this coupling is absent in AD mice.

### Intact PC Reactivation

Finally, we assessed reactivation of activity patterns within the PC. In NonTg mice, memory replay is compressed and preferentially occurs at 6x speed (Fig. 2A). So, we assessed template matches in 3xTg-AD at a range of compression factors to see if compressed template matching also occurred in 3xTg-AD mice (n=7 3xTg-AD data sets met the inclusion criteria described above for nonTg mice). Note, the lower number of 3xTg-AD data sets was due to fewer data sets with ≥6 tetrodes in PC (**Supplementary Table 1**). For 3xTg-AD mice, template matches oscillated around chance levels for all compression factors tested with two peaks, one at 6x and a second at 10x (Fig. 6, *Left*). First, we used a 6x compression factor, since this was the ideal compression factor for NonTg mice. We compared proportion of strong template matches during SWS between *pre-task-sleep* and *post-task-sleep* for both groups of mice. NonTg mice showed an increase in proportion of strong template matches in *post-task-sleep* relative to *pre-task-sleep* in 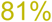 of data sets, while in 3xTg-AD mice, the proportion was 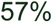, closer to chance levels but not significantly less than NonTg mice (**Fig**. **6**, *Middle*; χ^2^_(1)_ =2.1, p=0.15). While NonTg mice showed the strongest template matching when a 6x compression factor was applied, 3xTg-AD mice had weaker but equivalent template matching when a 6x or 10x compression factor was applied. Thus, we performed the same comparison as previously described for 6x compression between NonTg mice and 3xTg-AD mice with 10x compression applied. There was again no significant difference between the two groups, with both groups showing an increase in template matching in a moderate proportion of data sets (64% for NonTg and 57% for 3xTg-AD mice; **Supplementary Fig**. **2;** χ^2^_(1)_ =0.10, p=0.75). Thus, 3xTg-AD mice had a modest but non-significant reduction in template matching in PC regardless of compression factor 6x or 10x. In addition, to ensure that 3xTg-AD animals were not merely showing a change in the ideal compression factor, we compared data from NonTg mice with 6x compression factor to 3xTg-AD mice with 10x compression factor. The result was identical to the comparison made with a 6x compression factor applied for both genotypes (Fig. 6 *Middle*), suggesting that a shift in the rate of replay is not likely to explain the lower proportion observed in 3xTg-AD mice (81% for NonTg and 57% for 3xTg-AD mice; χ^2^_(1)_ =2.10, p=0.15). Finally, we assessed the strength of 6x compressed template matching (mean±SEM) centered on the DWT, a marker for PC memory replay. Template matches were z-scored and averaged for both pre-task-rest and post-task-rest for both groups. Both groups exhibited a decrease in template matches during the down state of the DWT, followed by an increase in matches during the upstate transition, with no significant difference between groups (Fig. 6 *Right*). Thus, the temporal relationship between template matches and DWTs remained consistent between both sleep sessions for both groups.

**Fig. 6.**
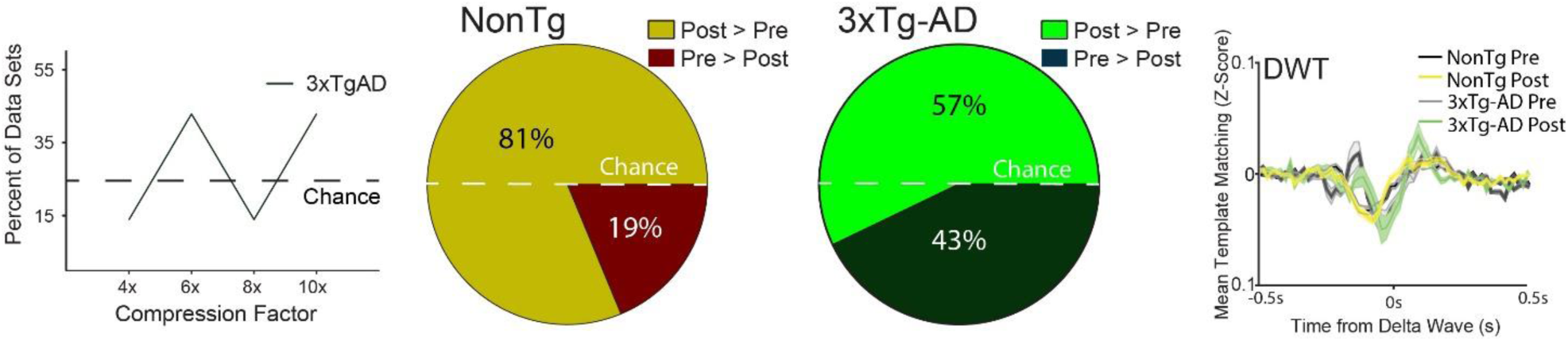
Modest changes in template matching in the parietal cortex (PC) of 3xTg-AD mice. *Left.* The percent of data sets for which template matches are greater in *post-task-sleep* and also greater than non-compressed data for 3xTg-AD mice indicates template matches oscillates around chance levels for all compression factors examined. *Middle.* The proportion of data sets in which there was a higher density of template matches in *post-task-sleep* for NonTg mouse data sets 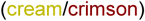 and 3xTg-AD mouse data sets (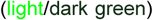/dark green). In PC there is a non-significant reduction in template matching in 3xTg-AD mice (χ^2^_(1)_=2.1, p=0.15). *Right*. Event-triggered average template matching Z-score (mean ± SEM) centered on the delta wave trough (DWT) for post-task-sleep for NonTg control 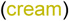 and 3xTg-AD 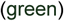 mice, relative to pre-task-sleep (black-NonTg/grey-3xTg-AD).

## Discussion

We set out to look for memory replay as a potential mechanism for a deficit in spatial reorientation learning that we had reported previously. We found that AD mice have increased sleep efficiency, but reduced SWR density. However, the longer duration of SWS compensated for reduced SWR density so that the total number of SWRs during rest periods was not reduced. Conversely, the density of DWs was not different for AD mice and as a result the total number of DWs was increased. The increased number of DWs may have partially ameliorated changes in memory reactivation with PC, because there was a non-significant reduction in *post-task-sleep* template matching in PC in AD mice. Finally, we found that in NonTg mice hippocampal SWRs slightly preceded PC DWs. This correlation was strengthened in *post-task-sleep*, and SWR-DW coupling was significantly correlated with performance on the *spatial reorientation task* the following day. However, in AD mice, SWR-DW coupling was significantly reduced and was not correlated with behavioral performance the following day. This suggests a decoupling between memory replay events for 3xTg-AD mice and subsequent behavioral performance, i.e., the expected benefit from memory reactivation during the intervening period between the behavioral training and testing is not occurring.

We observed a compensatory sleep pattern in our 3xTg-AD mice that were at early stages of disease progression (i.e., intracellular Tau and Aβ only, no plaques or tangles). Specifically, we found that AD mice had increased sleep efficiency, which increased the total number of SWRs, compensating for a reduction in density of SWRs. In addition, we observed that the increased sleep time also results in increased number of DWTs above the number observed in NonTg mice. Thus, the pattern of data we observed suggests it is possible that the increased sleep efficiency we observed is an early compensatory mechanism for changes in sleep physiology in the AD mice. Similar compensation have not been reported in humans; however, it is possible that sleep in humans that will develop AD has not been assessed at a parallel point in the disease progression. In humans, there are sleep changes, including a decrease in quality (increased fragmentation, increased time to sleep onset but retention of total time spent in sleep), and these changes begin to emerge in parallel with initial Aβ increase in the cerebrospinal fluid (Holth et al., 2017; Ju et al., 2013b). There is evidence for a bidirectional relationship between sleep and AD pathology. For example, a lack of sleep, as well as increased sleep disturbances have been associated with increased risk of AD, amyloid deposition, and cognitive impairment (Cordone et al., 2019; Liguori et al., 2017; Mander et al., 2017; Winer et al., 2019) and conversely there is evidence that SWS increases Aβ clearance (Ju et al., 2013a; Mander et al., 2016). Given that SWS clears Aβ, the increased sleep time we see prior to Aβ plaque formation may be a compensatory mechanism to dampen early pathological processes.

Like humans, animal models display sleep abnormalities concurrent with Aβ deposition (Colas et al., 2004; Huitrón-Reséndiz et al., 2002; Jyoti et al., 2010; Roh et al., 2012b; Schneider et al., 2014; Wisor et al., 2005; Zhang et al., 2005). Some mouse models show decreases in sleep after Aβ plaques begin forming (Huitrón-Reséndiz et al., 2002; Roh et al., 2012a; Van Erum et al., 2019), suggesting that the increased sleep we observed may be an early compensatory mechanism that eventually breaks down as pathology progresses. In contrast to what we observed in 3xTg-AD mice, an amyloidosis only mouse model (APP/PSEN) had decreased in time spent in sleep prior to the development of plaques (Jyoti et al., 2010). Later in disease progression, after the development of Aβ plaques, a different Aβ mouse model, Tg2576, did not have deficits in sleep time (Wisor et al., 2005), suggesting the effects of Aβ alone might produce a different pattern of sleep changes over the course of disease progression than we observed in 3xTg-AD mice which have both Tau and Aβ pathology. Tau alone also influences sleep; post-tangle formation PLB2_Tau_ mice have reduced total time spent in sleep, as well as NREM sleep (Koss et al., 2016). Thus, the pre-plaque and tangle effects we observed in the 3xTg-AD mouse model may be the result of either early tau pathology, or the interaction between tau and Aβ. Together, this past and present data suggests that the influence of tau and Aβ on sleep varies over the course of disease progression and that interactions between tau and Aβ may produce a different phenotype than either pathology component alone.

Abnormally rigid sequences of cell activity have been reported in AD mice in the mature stages of disease progression, after neurodegeneration (Cheng and Ji, 2013; Gillespie et al., 2016; Witton et al., 2016). Mouse models of Tauopathy and familial AD have shown changes in characteristics of hippocampal sharp wave ripples (SWRs), after neurodegeneration (Cheng and Ji, 2013; Gillespie et al., 2016; Witton et al., 2016). These changes include decreased density of SWRs, consistent with our results (Ciupek et al., 2015; Gillespie et al., 2016). However, these reports did not assess the influence of sleep on total numbers of SWRs. Our findings suggest that deficits in the density of SWR generation may appear early in AD, but at least initially, compensatory sleep changes may prevent a reduction in the total number of SWRs during sleep.

Replay of activity patterns during sleep is critical for learning and memory, including navigation (Jadhav et al., 2012; Staresina et al., 2013). The hippocampal formation is crucial to the storage of ‘episodic’ memories (memories for experiences that unfold in space and time), and for assisting the neocortex to extract generalized knowledge from these specific experiences (Eichenbaum, 2010). Because the hippocampus is critical for recent memories, it has been suggested that it generates a unique code reflecting the spatial or temporal context of experience (Buzsáki, 1989; Marshall and Born, 2007; Nadel et al., 2000; Teyler and DiScenna, 1986). This code has been suggested to provide a tag or ‘index’ that links together components of a given experience that are independently stored in weakly interacting modules throughout the neocortex, including the PC (Burke et al., 2005; Skelin et al., 2018). The hippocampal output at the time of the experience has been theorized to enable indirect, coordinated retrieval of episodic information from these modules. For recent memories, the hippocampus appears to orchestrate retrieval of information throughout the cortex (Siapas and Wilson, 1998). The process may be initiated in the neocortex for older memories (Kubie et al., 1999; but see Nadel et al., 2000; Scoville and Milner, 1957); however, more recent theories and data suggest that hippocampal involvement may remain much longer than originally theorized (Kubie et al., 1999; Nadel et al., 2000). We found that in 3xTg-AD mice that have not yet presented with tangles and plaques, there is a robust deficit in hippocampal-PC interactions during sleep, while memory replay events in PC are not significantly altered. Thus, the learning and memory deficits we observed may represent a failure of memory indexing or trace formation as a result of altered cotrical-hippocampal interactions. In the 3xTg-AD mouse model, we see compensatory sleep changes early in disease progression, with increased sleep time compensating for decreases in SWR count and giving rise to an increase in DWT number. However, these changes fail to compensate for all dysfunction. Impairments remain in hippocampal-PC interactions, with deficits in experience induced increases in SWR by DWT interactions. Furthermore, these hippocampal-PC deficits correlate with deficits in performance on the virtual maze. Thus, AD may cause hippocampal-cortical network changes which impair spatial orientation because of impaired learning related plasticity particularly between PC and hippocampus during SWS.

## Acknowledgements

This research was supported by grants from NIA AG049090 to AAW and B.L.M. was supported through DARPA HR0011-18-2-0021, NSERC Grant RGPIN-2017-03857 and Canadian Institutes of Health Research (CIHR) grant PJT 156040.

## Notes

**Conflict of Interest:** The authors declare no competing financial interests.

