## Supplementary Table 1 for "Impaired Hippocampal-cortical interactions during sleep and memory reactivation without consolidation in a mouse model of Alzheimer’s disease"

| | Technical Issues (e.g. tracking problems) | > 9 trials | Still ≥ 20 minutes per sleep session | ≥ 6 tetrodes in PPC* | Total included (cross correlation, PPC reactivation) |
|---|---|---|---|---|---|
| NonTg | 0 | 8 | 6 | 2 | 16, 14 |
| 3xTg-AD | 4 | 6 | 7 | 6 | 11, 7 |

**Supplementary Table 1. Number of excluded data sets by genotype.** The number of data sets excluded for each exclusion criteria is shown for each genotype. *only applied to cross correlation analysis.
