## Supplementary figure 1 for "Impaired Hippocampal-cortical interactions during sleep and memory reactivation without consolidation in a mouse model of Alzheimer’s disease"

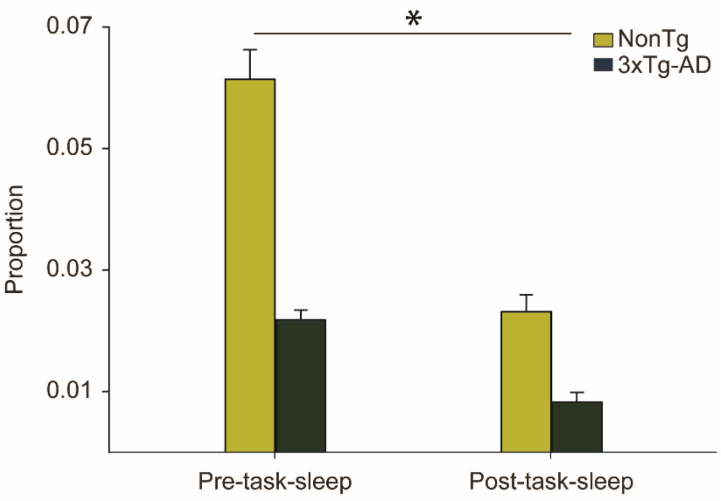


**Supplementary Figure 1. REM Sleep is reduced following spatial reorientation training.** REM sleep as a proportion of total sleep time for 3xTg-AD (cream) and NonTg mice (dark green). There was a significant reduction in the proportion of REM sleep between *pre-task-sleep* and *post-task-sleep* (sleep phase: $F_{(1, 25)}$=4.67, $p<0.05$). There was not an effect of group or group by sleep session interaction ($Fs_{(1, 25)}$<2.5, ps>0.13).
