## Supplementary figure 2 for "Impaired Hippocampal-cortical interactions during sleep and memory reactivation without consolidation in a mouse model of Alzheimer’s disease"

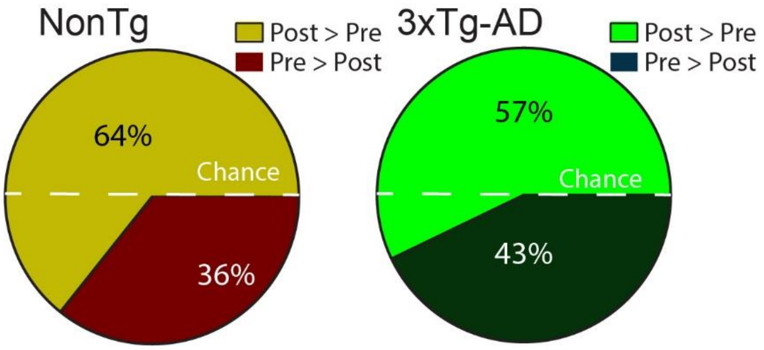


**Supplementary Fig. 2 Template matching in parietal cortex (PC) is unchanged in AD mice butr reduced in NonTg mice when a 10x compression factor is applied.** The proportion of data sets in which there was a higher density of template matches in *post-task-sleep* for NonTg mouse data sets (cream/crimson) and all 3xTg-AD mouse data sets (light/dark green). In PC there is no difference in template matching in 3xTg-AD versus NonTg mice when a 10x compression factor is applied ($\chi^2_{(1)}$=0.10, p=0.75). The proportion of data sets with stronger post-task-sleep template matching is slightly reduced in NonTg mice from 75% to 64% and remains the same in 3xTg-AD mice 57%.
